# The Loroxanthin cycle: A new type of xanthophyll cycle in green algae (Chlorophyta)

**DOI:** 10.1101/2021.10.28.466245

**Authors:** T E van den Berg, R. Croce

**Author notes:** Corresponding author: R. Croce. Integrated devices and systems group, University of Twente Drienerlolaan 5, 7522 NB, Enschede, The Netherlands.

## Abstract

Xanthophyll cycles have proven to be major contributors to photoacclimation for many organisms. This work describes a light-driven xanthophyll cycle operating in the chlorophyte *Chlamydomonas reinhardtii* and involving the xanthophylls Lutein (L) and Loroxanthin (Lo). Pigments were quantified during a switch from high to low light and at different time points from cells grown in Day/night cycle. Trimeric LHCII was purified from cells acclimated to high or low light and their pigment content and spectroscopic properties were characterized. The Lo/(L+Lo) ratio in the cells varies by a factor of 10 between cells grown in low or high light leading to a change in the Lo/(L+Lo) ratio in trimeric LHCII from 0.5 in low light to 0.07 in high light. Trimeric LhcbMs binding Loroxanthin have 5±1% higher excitation energy transfer from carotenoid to Chlorophyll as well as higher thermo- and photostability than trimeric LhcbMs that only bind Lutein. The Loroxanthin cycle operates on long time scales (hours to days) and likely evolved as a shade adaptation. It has many similarities with the Lutein-epoxide - Lutein cycle of plants.

## Introduction

Green algae are found worldwide in a large variety of habitats: from the dessert crust (Perera et al., 2018) to the pole-ice (Kirst and Wiencke, 1995). Their ability to grow under different conditions follows from the millions of years of evolution after their ancestor incapsulated a cyanobacterium. Their evolutionary success did not rely on the photosynthetic electron transfer chain that has remained largely unchanged but instead depended on the versatile acclimation machinery that evolved around these fundamental processes (Ballottari et al., 2012; Leliaert et al., 2012; de Vries and Archibald, 2018).

In all organisms performing oxygenic photosynthesis, light-harvesting and trapping of excitation energy (EE) occur in photosystems (PS) I and II. The absorption cross-section of the PS core complexes is extended by an outer antenna, which in plants and green algae is composed of members of the light-harvesting multigenic family (Croce and van Amerongen, 2020; Pan et al., 2020). LHCII is the main antenna complex in plants and green algae, it can be associated with both photosystems and is mainly present in trimeric form. In the *Chlorophyte C. reinhardtii*, LHCII is composed of nine LHCBM proteins (Minagawa and Takahashi, 2004; Natali and Croce, 2015). In addition to LHCII, *C. reinhardtii* contains PSI-specific antennae, called Lhca (Mozzo et al., 2010) and the monomeric antennae CP26 and CP29, mainly associated with PSII (Elrad et al., 2002). A large variety of carotenoids is present in plants and green algae (Takaichi, 2011; Jeffrey et al., 2017) acting as light sensors, shading pigments, antioxidants, membrane stabilizers, light-harvesting pigments and excitation-energy quenchers (Satange, 2016). A selection of carotenoids active in light-harvesting and photoprotection is associated with photosynthetic proteins. β-carotene is mainly bound to the PS I and II core, while xanthophylls are bound to the light-harvesting complexes of plants and green algae (Caffarri et al., 2014).

Four carotenoid binding sites are present in LHCII, that are highly conserved in plants and green algae (Fig. 1). The N1 site is highly specific for Neoxanthin in most complexes. The L1 site is usually occupied by Lutein and the L2 site also preferentially binds Lutein (Natali and Croce, 2015; Pan et al., 2020). Studies on the LHCII of plant showed that the xanthophylls in all three bindings sites are involved in light-harvesting (Croce et al., 2000), while only L1 and L2 are responsible for Chl^3^ quenching (Mozzo et al., 2008). The fourth carotenoid binding site, V1, is located at the periphery of the complex and is occupied either by Violaxanthin or Lutein. Experiments on plants have shown that in this pocket, the carotenoid is only weakly bound to the complex, not involved in light-harvesting and is easily lost during the isolation of the complex from the membrane (Caffarri et al., 2001).

**Figure 1.**
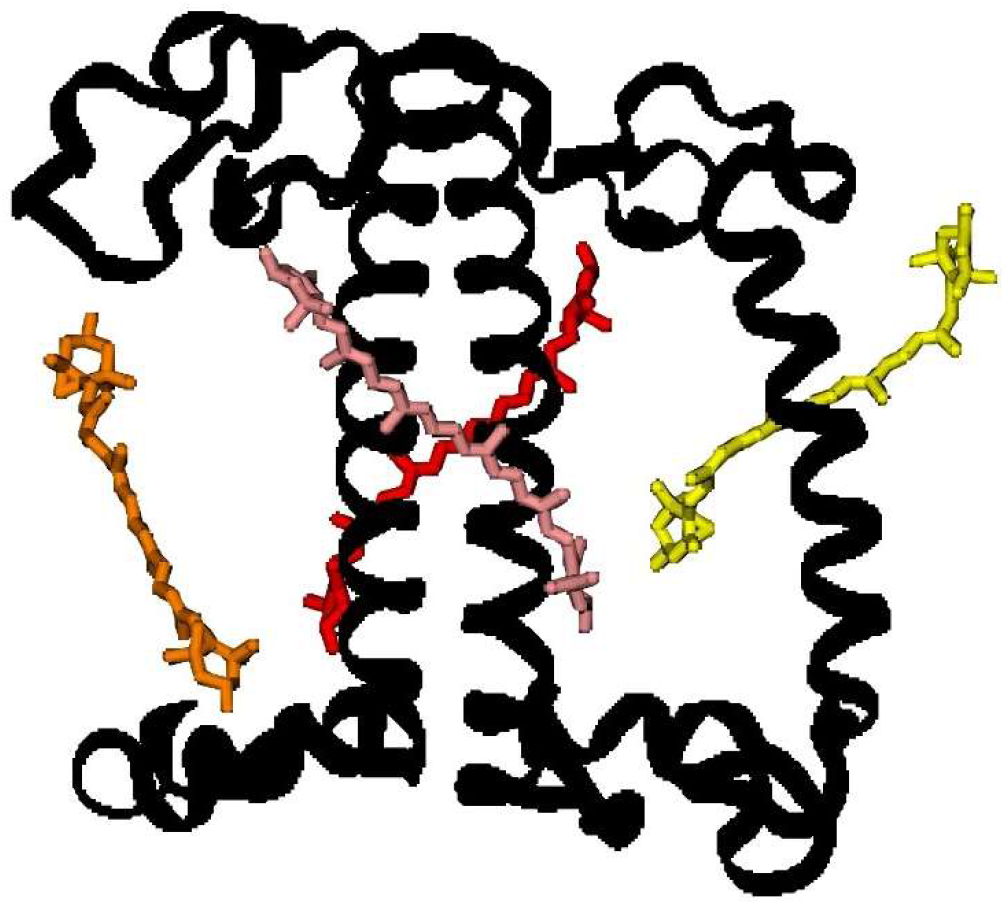
The carotenoid binding sites of LHCII monomer (from (Shen et al., 2019)) Neoxanthin in the N1 site (yellow),Lutein in the L1 (red) and L2 (pink) sites and Violaxanthin in the V1 site (orange) were modeled according to the structure of plant LHCII (Liu et al., 2004). For clarity, the chlorophylls are omitted.

The light-harvesting antennae are involved in multiple light-acclimation processes. These are vital for plants and green algae because they allow the fine-tuning of photosynthesis, preventing photodamage or photo-starvation. The acclimation processes occur on different time-scales as responses to changes in light quality and quantity (Wobbe et al., 2016): from short-term processes that act on seconds to minutes to long-term processes lasting for hours to days. Long-term processes are connected to regulated changes in gene expression that affect the capacity of photosynthetic electron transport, respiration and light-harvesting (Falkowski and Chen, 2003), CO_2_ concentrating mechanisms and CO_2_ fixation (Giordano et al., 2005), as well as the capacity for photoprotection and shading (Wobbe et al., 2016). Short-term processes include non-photochemical quenching (NPQ), a process that reduces the excited-state lifetime of the chlorophylls (Chl), thus preventing ^3^Chl formation that can lead to the production of damaging reactive oxygen species (ROS) (Bassi and Dall’Osto, 2021). NPQ is activated by the acidification of the lumen. In *Chlamydomonas reinhardtii*, several proteins are involved in the fast phase of NPQ called qE (s to min): LHCSR1 and LHCSR3 (Peers et al., 2009; Dinc et al., 2016) and PsbS (Correa-Galvis et al., 2016; Tibiletti et al., 2016). In addition, some algae contain the so-called Xanthophyll cycles (XC), which have multiple roles in photoprotection (García-Plazaola et al., 2007). Thus far, six such cycles are known (García-Plazaola et al., 2007). The most extensively studied are the Violaxanthin - Antheraxanthin - Zeaxanthin cycle (VAZ) present in plants and green algae, the Diadinoxanthin - Diatoxanthin cycle (Ddx) present in diatoms and the Lutein-epoxide – Lutein cycle (LLx) present in some plants. Out of the other three cycles two are truncated versions of the VAZ cycle, namely the Violaxanthin - Antheraxanthin cycle (VA) (Goss et al., 1998; Stamenković et al., 2014) and the Antheraxanthin – Zeaxanthin (AZ) cycle (Rmiki et al., 1996). The final one is the Lutein – Siphonaxanthin (LS) cycle (Raniello et al., 2006) (for more details see (García-Plazaola et al., 2007). Most cycles consist of the de-epoxidation of xanthophylls in (light) stress and their epoxidation in the absence of stress (low light or darkness) (Goss and Latowski, 2020; Fernández-Marín et al., 2021). The epoxy- and epoxy-free xanthophylls have different properties that favor light-harvesting or photoprotection (Havaux, 1998). The VAZ cycle and the Ddx cycle are well-known to contribute to NPQ (Goss and Jakob, 2010). A role in NPQ has also been suggested for the LLx cycle (Esteban et al., 2010; Matsubara et al., 2011; Leonelli et al., 2017). The activation of the xanthophyll-dependent NPQ requires several minutes and it is thus slower than the LHCSR/PSBS dependent qE, but still very fast compared to other acclimation processes (Stamenković et al., 2014; Quaas et al., 2015; Christa et al., 2017). In addition to their role in NPQ, the XCs protect from photoinhibition in high light by enhancing the antioxidant activity in the membrane (Havaux and Niyogi, 1999; Havaux et al., 2007; Johnson et al., 2007; Lepetit et al., 2010) and the membrane stability (Havaux, 1998; Gruszecki and Strzałka, 2005; Bojko et al., 2019). Upon a switch to low light the XCs provide advantages in light-harvesting by decreasing the energy losses through NPQ (VAZ) (Kromdijk et al., 2016) and increasing the carotenoid to chlorophyll excitation energy transfer (EET) (LLx) (Matsubara et al., 2007; Leonelli et al., 2017). All XCs are active upon a change of light intensity but the turnover kinetics can vary from minutes (VAZ, Ddx) to days (LLx) depending on the type of cycle (García-Plazaola et al., 2007; Goss and Jakob, 2010), the plant species (LLx) (García-Plazaola et al., 2007), temperature (VAZ) (Reinhold et al., 2008) and the changes in light intensity (VAZ) (Kress and Jahns, 2017). Additionally, the XC pool size can be highly variable, ranging from a minor fraction to the dominating xanthophylls (García-Plazaola et al., 2007; Stamenković et al., 2014) and not all xanthophylls in the pool may be available to the cycle (Snyder et al., 2005). Lastly, XCs might influence the composition of the free pool of xanthophylls in the membrane, the xanthophylls bound to the light-harvesting antenna or both, depending on the cycle, the species and the duration of the stress condition (Snyder et al., 2005; Matsubara et al., 2007; Reinhold et al., 2008; Xu et al., 2015).

The unicellular green alga *Chlamydomonas reinhardtii* became a model green alga because it grows quickly in the lab and is easy to cross (Sasso et al., 2018). Its PS and their antenna e.g. (Mozzo et al., 2010; Natali and Croce, 2015; Shen et al., 2019; Suga et al., 2019; Huang et al., 2021) and photoacclimation behavior (Bonente et al., 2012; Allorent et al., 2013; Nawrocki et al., 2016, 2020; Polukhina et al., 2016) are well characterized. Interestingly, although the VAZ XC was observed in this alga, this cycle is not involved in NPQ (Bonente et al., 2011; Quaas et al., 2015).

One of the xanthophylls of *C. reinhardtii* is Loroxanthin, which is formed by the hydroxylation of the methyl group at C9 of the polyene of Lutein by an unknown enzyme (Grossman et al., 2004). It is widespread among Chlorophyte, Euglenophyte and Chlorarachniophyte (Takaichi, 2011) and it binds to the light-harvesting complexes (Bishop, 1996; Mozzo et al., 2010; Natali and Croce, 2015; van den Berg et al., 2018). Loroxanthin is associated with the PSI and PSII supercomplexes of low light-grown *C. reinhardtii* (Pineau et al., 2001; Drop et al., 2011, 2014). Furthermore, Loroxanthin abundance and its ratio with Lutein vary in different light intensities in *C. reinhardtii* and other Chlorophyte (Bishop et al., 1989; Garrido et al., 2009; Bonente et al., 2012; van den Berg et al., 2019). We hypothesize that the differences in cellular Lutein and Loroxanthin content observed in low and high light grown cultures are due to a light-driven XC (Fig. **2a**, spectra in **2b**) that affects the Xanthophyll composition of the light-harvesting complexes and thereby their properties. In this work we tested this hypothesis using a combination of biochemical and spectroscopic analyses at the protein and cellular levels.

**Figure 2.**
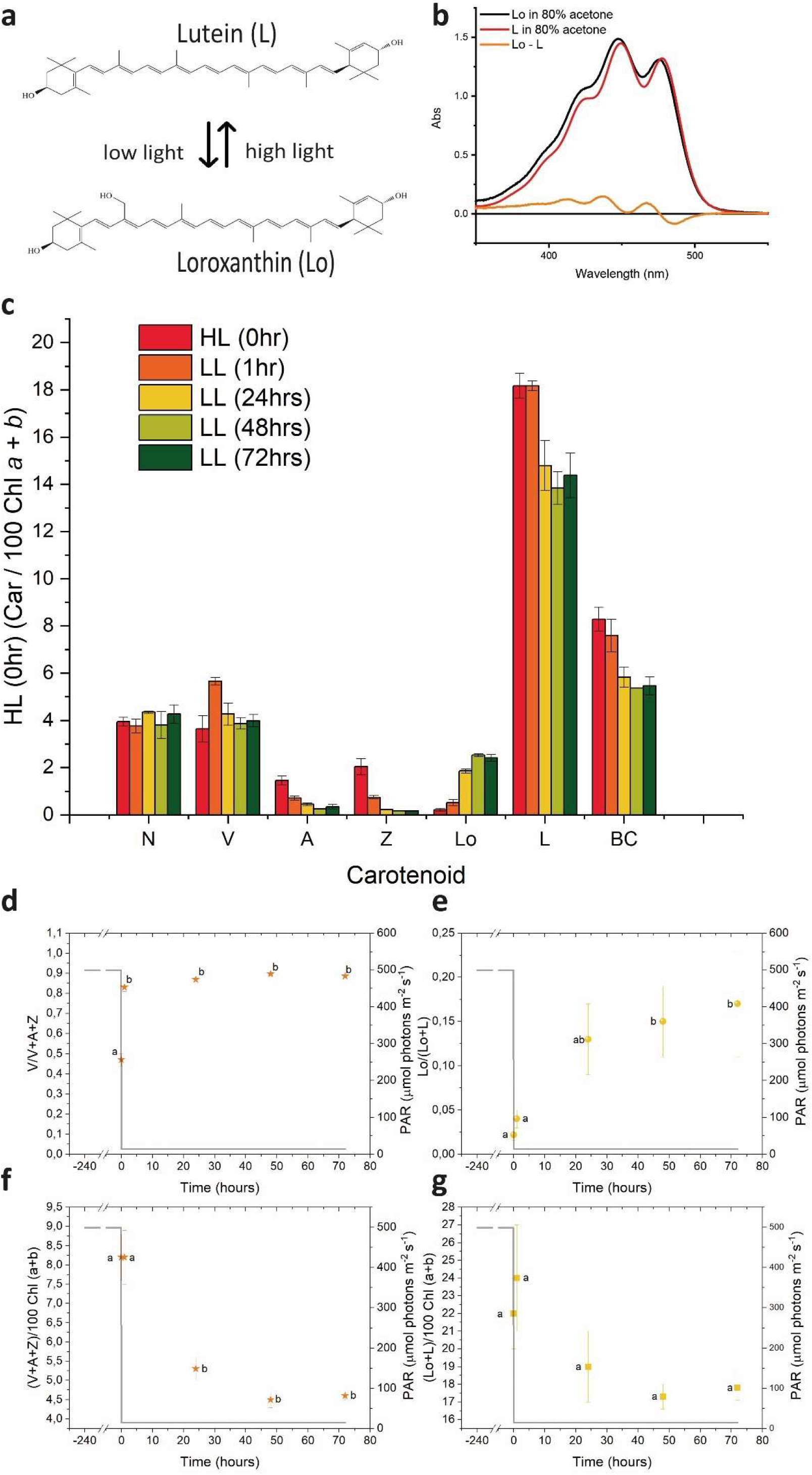
Spectra of Loroxanthin and Lutein and the changes in their content at different light intensities. **A** Chemical structures of Lutein and Loroxanthin and an illustration of their relationship at different light intensities. **B** Absorption spectra of Loroxanthin and Lutein in 80% acetone. The difference spectrum is also shown. **C** Carotenoid composition (relative to 100 Chl (*a*+*b*)), **D** VAZ epoxidation state, **E** LLo hydroxylation state and VAZ (**F**) as well as LLo (**G**) pool size (relative to 100 Chl (*a*+*b*)) of *C. reinhardtii* cells at different time points (0, 1, 24, 48 and 72hours) following the shift of the culture from continuous HL to continuous LL. Averages represent two technical replicas per five biological replicas per timepoint. Error bars represent the standard error. Letters (a,b,ab) indicate significant differences between groups (ANOVA). N, Neoxanthin; V, Violaxanthin; A, Antheraxanthin; Z, Zeaxanthin; Lo, Loroxanthin; L, Lutein; BC, β-carotene.

## Materials and methods

### Experimental design

To check the existence of a Lutein - Loroxanthin cycle in *C. reinhardtii*, we exposed the cells to changes in light intensity during growth and we measured the pigment content at different time points during light acclimation. The first set of experiments was performed on cells acclimated (>10 days) to continuous high light (HL) (500 µmol photons m^-2^s^-1^) shifted to continuous low light (LL) (15 µmol photons m^-2^s^-1^). In the second set of experiments, the cells were instead grown in a day/night cycle using (18h:6h day/night; D/N) a sinusoidal light regime (peak light intensity 1300 µmol photons m^-2^s^-1^).

### Growing conditions

*Chlamydomonas reinhardtii* CC-124 was grown photoautotrophically in high salt medium (HSM) (Sueoka, 1960) in low light (LL, 15 μmol m^-2^ s^-1^ white fluorescent bulbs) or high light (HL, 500 μmol m^-2^ s^-1^ white fluorescent bulbs) shaking at 150 rpm at room temperature or it was grown in a DN cycle (18D:6N, sinusoidal of warm white LED light, 1300 μmol m^-2^ s^-1^ at midday (hr 9)) in a bubble column photobioreactor (PSI, Czech republic) at 25 °C. Biological replicates consist of independent cultures that were acclimated for at least 10 generations at each culturing condition (D/N or prior to transfer to a different light intensity) with culture dilution every 2-3 days, maintaining the cells in the exponential growth phase.

### Purification of antenna complexes

Thylakoids were prepared as previously described (Chua and Bennoun, 1975) with the addition of 1 mM benzamidine and ε-aminocaproic acid to the buffer (Tricine NaOH pH 7.8 instead of Hepes pH 7.5). In short, the thylakoid membranes were purified using a discontinuous sucrose gradient (100000xg, 1 h, 4°C in a SW41 swinging bucket rotor). Thylakoids were pelleted, unstacked with 5mM EDTA, washed with 10mM tricine-NaOH (pH 7.8) and homogenized in the solubilization buffer (10 mM Tricine-NaOH, 150 mM NaCl, pH 7.8 pH. Subsequently, thylakoids (kept in the dark on ice) diluted at a Chl concentration of 1.0 mg mL^−1^ were mixed with an equal volume of solubilization buffer containing freshly prepared detergent (0,6 % Dodecyl-α-d-maltoside (α-DM) (Anatrace)). The thylakoids were solubilized for 20 min at 4 °C in the dark with end-over-end shaking. Isolation of complexes was performed by sucrose density gradient made by freezing and thawing 0.5M sucrose, 10 mM Tricine-NaOH, 0,05 % α-DM (pH 7.8) centrifuged at 240000xg for 17 hours at 4 °C in a SW41 rotor. Green bands were collected with a syringe, flash-frozen in liquid nitrogen and stored at -80 °C until use.

### Steady-state absorption, fluorescence, and circular dichroism

The sample buffer used for all room temperature (RT) experiments was 0.5 M sucrose, 20 mM Tricine (pH 7.8), 0.05% alpha-DM. 66% (w/w) glycerol was added to the buffer for 77 K experiments. Sample OD at the maximum in the Qy region was 0.8–1 for absorption and circular dichroism (CD) and below 0.05 for fluorescence measurements. RT and 77 K absorption spectra were recorded with a Cary 4000 spectrophotometer (Varian) with a spectral bandwidth of 2 nm. For 77 K measurements samples were cooled in a cryostat (Oxford Instruments). 77K absorption spectra were measured with a UV-2600 spectrophotometer (Shimadzu) with a spectral bandwidth of 2 nm. Fluorescence emission spectra at RT and 77 K were recorded on a Fluorlog 3.22 spectrofluorimeter (Jobin-Yvon spex). For fluorescence emission spectra, the spectral bandwidths were 3 nm for excitation (440, 475 and 500 nm), and 1 nm for emission. Excitation spectra were recorded at 735 nm emission with the spectral bandwidths 1 nm for excitation, and 3 nm for emission. 735 nm was chosen in order to record the excitation spectra up to 700 nm. Excitation spectra at 680 nm and 735 nm were identical. An optical filter was placed before the detector to block light <600 nm for emission or <700nm for excitation spectra. CD spectra were recorded at 20 °C with a Chirascan CD spectrophotometer (Applied Photophysics) equipped with a temperature control unit TC125 (Quantum Northwest). The spectral bandwidth was 1 nm and the sample volume 400 µL. The thermostability of the complexes was determined by the loss of the CD signal between 450 and 550 nm as a function of increasing temperature (20-90 °C, 2.5 °C steps). Each heating step took 1 min, followed by 1 min of equilibration before measuring the spectrum.

### Pigment analyses

The pigment composition was determined by fitting the absorption spectrum of the 80% acetone extracts with the spectra of the individual pigments in the same solvent as earlier described (Croce et al., 2002) and by HPLC. HPLC was performed on a System gold 126 equipped with a 168 UV–VIS detector (Beckman Coulter, USA) using a C18-Sphereclone column (Phenomenex 5U ODS1, 00G-4143-E0, 4.6 mm × 250 mm). Loroxanthin, Neoxanthin, Violaxanthin and Chl *b* were separated according to an earlier described protocol (Pineau et al., 2001). Because with this method the separation between Chl *a* and Lutein could not be achieved on our column, Violaxanthin, Antheraxanthin, Zeaxanthin, Lutein, Chl *b*, Chl *a* and β-carotene were additionally separated using another protocol (van den Berg et al., 2019).

### Time-resolved fluorescence

Time-resolved fluorescence was measured at RT by a time-correlated single-photon counting (TCSPC) setup (FluoTime 200 fluorometer, PicoQuant). Samples were stirred with a magnetic bar in a 1 cm quartz cuvet. Excitation was performed with a laser diode at 438 nm, with 5 MHz repetition rate and 1 µW power. Careful checks at higher and lower power confirmed the absence of non-linear processes (e.g., annihilation). Fluorescence was detected with 4 ps timesteps, at 680 nm (8 nm bandwidth), at an angle of 90° with the excitation, through a polarizer set at the magic angle relative to the excitation polarization. The instrument response function (FWHM 88 ps) was determined using pinacyanol iodide in methanol (6 ps lifetime (Van Oort et al., 2008)). Data were accumulated until the number of counts in the peak channel was 20,000. Fluorescence decay curves were fitted with a multi-exponential decay, with amplitudes and lifetimes convoluted with the IRF with the Fluofit software (Pico-Quant). Three components were necessary to get a good fit of the data as judged by χ2, the distribution of the residuals around 0 and the autocorrelation function of the residuals.

### Carotenoid EET efficiency

Energy transfer efficiencies from Car-to-Chl a were estimated by fitting the fluorescence excitation spectra and the 1-T spectra with the spectra of the individual pigments and comparing the contribution of the same pigment to the two spectra. The 1-T and the excitation spectrum were normalized to the fitted quantity of Chl *a* (100% efficiency of Chl *a* - > Chl *a* EET). Deconvolution of spectra in the 400-520 nm wavelength range was performed as described in (Croce et al., 2000).

### LHC photostability

The photostability of the complexes was determined by following the loss of absorption between 350 and 750 nm as a function of illumination time. The OD of the sample at 435nm was 0.45 and the volume 500 µL. Samples were measured in a quartz cuvette with 1 cm path length. Samples were illuminated with white light from a halogen lamp (4500 μmol m^-2^ s^-1^) equipped with an optical fiber arm through a 1 cm plastic cuvette filled with water and cooled by a fan to minimize heating by the light source during illumination. After each illumination time, the sample was mixed and an absorption spectrum was recorded.

#### Statistical Test

Means were compared by paired, double-sided, students t-test, with the pairs representing the timepoints of individual biological replicas or in the case of LHCII, the individual preparations.

## Results

### Changes in Lutein and Loroxanthin content upon transfer from High light to Low light

Pigment analysis showed that the Lo/(L+Lo) ratio (Lutein hydroxylation state) increased ∼eight times in the 72 hours following the transfer from HL to LL (Fig 2E). In addition, the total L+Lo pool size (normalized to the total Chl content, Fig 2G) decreased by 19% upon transfer to LL. The VAZ XC also showed changes during the experiment. The VAZ epoxidation state increased ∼two times in the 72 hours after the transfer from HL to LL (Fig 2D) and the total VAZ pool size was reduced by 44% (Fig. 2F). The Chl concentration of the cultures transferred to LL increased 2.8 ± 0.5 times (Supporting information Fig. S1).The Chl/Car ratio also increased from 2 to 2.6 ± 0.2 in 72 hours LL, while the Chl *a*/*b* ratio was unaffected (Supporting information Fig. S1).

### Lutein-Loroxanthin changes during an 18:6 (D/N) sinusoidal light regime

To check if the changes in Lutein and Loroxanthin content also occurred in light conditions mimicking the natural environment, we monitored the levels of all xanthophylls during a simulated summer day (18:6 D/N cycle) (Fig. **3(A)**). The Lo/(L+Lo) ratio varies throughout the day and roughly follows the inverse of the light intensity (0-1300 μmol m^-2^ s^-1^) (Fig. **3(C)**). The Lo/(L+Lo) ratio at any time during the D/N cycle (Fig. **3(C)**) was much smaller than in the cultures shifted (>24 hours) from HL to LL (Fig **2(E)**). The minimum of the Lo/(L+Lo) ratio was reached after 9-12 hours of light and was delayed with respect to the de-epoxidation state of the VAZ cycle (max 6-9 hours) (Fig. **3(B**,**C)**). Chl *a*/*b* increased from 2.5 ± 0.1 to 2.8 ± 0.1 and Chl/Car decreased from 2.1 ± 0.1 to 1.9 ± 0.1 in the first 12 hours of light and returned to the starting values at the end of the light period (Supporting information Fig. S2).

**Figure 3.**
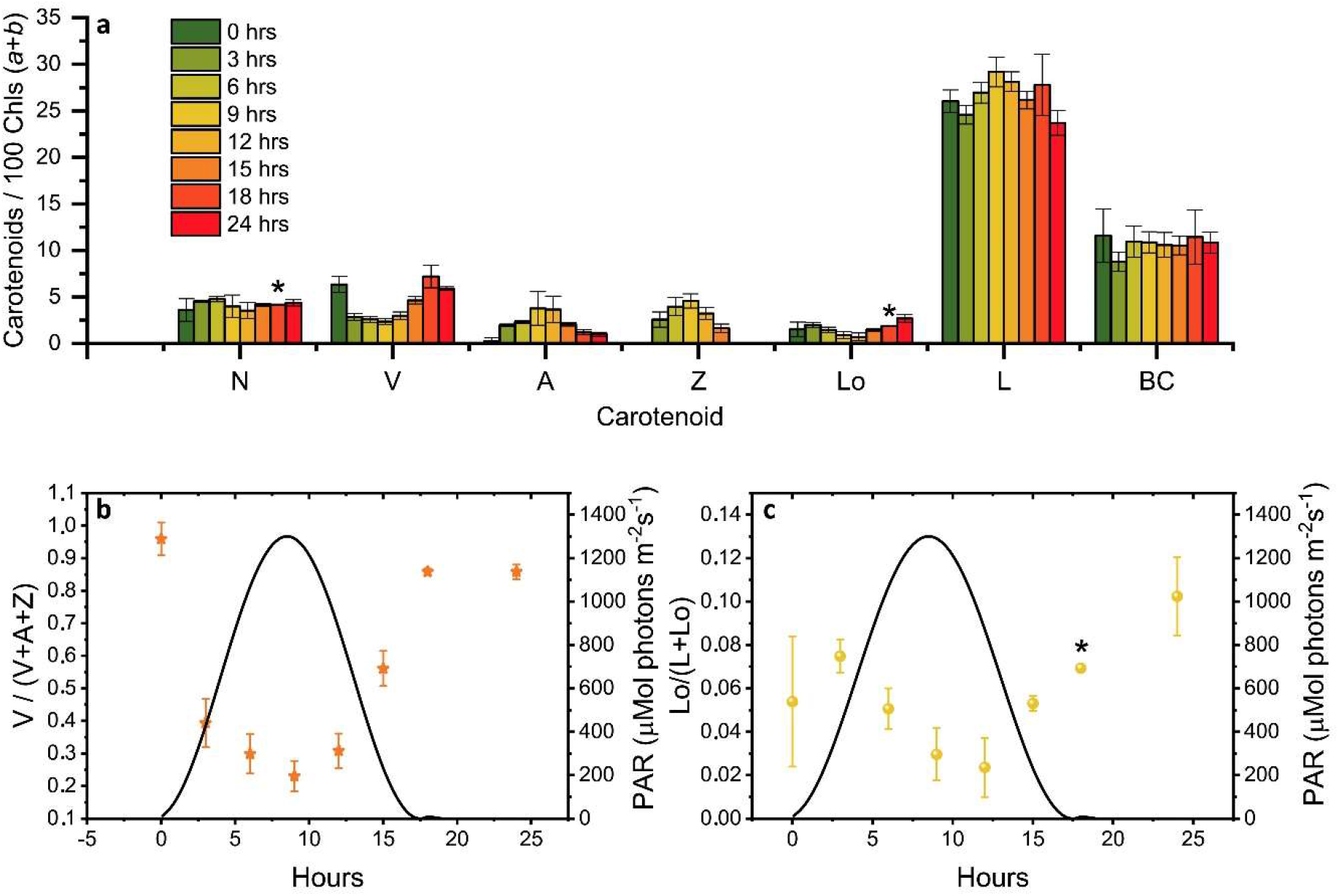
Changes in the carotenoid composition of *C. reinhardtii* cells during a “simulated summer” day (18:6 (D:N) sinusoidal light regime). **A** Carotenoid composition relative to 100 Chl (*a*+*b*) molecules of cultures at different time points (0, 3, 6, 9, 12, 15, 18 and 24 hours). The sample at “0 hours” was taken right before dawn. The results are the mean of three biological replicas. The error bar represents the standard error of the mean. **B** Epoxidation state of the VAZ pool. **C** Hydroxylation state of the L-Lo pool. N, Neoxanthin; V, Violaxanthin; A, Antheraxanthin; Z, Zeaxanthin; Lo, Loroxanthin; L, Lutein; BC, β-carotene. Full statistics are shown in SM Table 1. * indicate that data from one biological replica is missing due to loss of sample (t=18).

In summary, the cellular content of Lutein and Loroxanthin is affected by changes in light intensity. The changes in Lo/(L+Lo) ratio can be substantial when cells are adapted for days to different light conditions, but they are slow relative to the changes in the VAZ cycle.

### Lutein and Loroxanthin content in trimeric LHCII of high- or low-light acclimated C. reinhardtii

To investigate if the light-induced changes in Lutein and Loroxanthin content affect the pigment composition of LHCII, we purified trimeric LHCII from cultures that were photo-acclimated to HL (LHCII-HL) or LL (LHCII-LL). The complexes were isolated by mildly solubilizing the thylakoid membranes and loading them on a sucrose density gradient. The pigment composition (Table 1) shows that while LHCII-LL binds a similar amount of Loroxanthin and Lutein (1.3 per monomer), LHCII-HL binds mainly Lutein (2.5 molecules per monomer) and contains only traces of Loroxanthin. The 86% decrease of Loroxanthin in LHCII-HL compared to LHCII-LL is compensated by an increase of Lutein. Other differences in pigment composition were a slightly higher Chl (*a*/*b*) ratio and a decrease of V in LHCII-HL compared to LHCII-LL.

**Table 1.** **Pigment composition of LHCII-HL and LHCII-LL. Xanthophyll content is normalized to 14 Chls (*a*+*b*) molecules to indicate the xanthophylls per monomer in the trimeric complex. Chl *a*/*b* and Chl/Car are expressed in mol/mol. n indicates the number of biological replicas. * The means are significantly different P < 0.05**.

| *LHCII* | *Chl*<br>*a/b* | *Chl/*<br>*Car* | *N* | *V* | *A* | *Lo* | *L* |
| --- | --- | --- | --- | --- | --- | --- | --- |
| *LHCII-LL*<br>(*SE*, *n*=4) | 1.17*<br>(0.04) | 3.74<br>(0.18) | 0.76<br>(0.03) | 0.49*<br>(0.04) | 0 | 1.3*<br>(0.3) | 1.3*<br>(0.2) |
| *LHCII-HL*<br>(*SE*, *n*=4) | 1.29*<br>(0.05) | 3.78<br>(0.13) | 0.6<br>(0.1) | 0.25*<br>(0.03) | 0.08<br>(0.13) | 0.18*<br>(0.06) | 2.5*<br>(0.1) |

### Spectral characteristics of trimeric LHCII binding Lutein or Loroxanthin

The absorption (**A**,**B**), circular dichroism (CD) (**C**) and fluorescence (**E**,**F**) spectra of LHCII-HL and LHCII-LL (Fig. **4**) display very similar characteristics. The small differences in absorption around ∼475 and ∼650 nm for LHCII-LL compared with LHCII-HL reflect the difference in Chl*a*/*b* ratio. The lower absorption at ∼674 nm and higher at ∼665 nm and ∼680 nm at 77K for LHCII-LL compared with LHCII-HL indicate small differences in the energy of some Chls *a* (Fig 4(**A**,**B**)). The higher absorption at ∼513 nm and lower at ∼497 nm for LHCII-LL compared with LHCII-HL, reflect changes in the carotenoid composition. This also explains the difference in the CD spectra in the 440-500 nm range (Fig. **4(C)**). The second derivative of the absorption spectra measured at 77K (Fig. **4(D)**) demonstrates that both LHCII-LL and LHCII-HL have their lowest energy carotenoid transition around ∼510 nm, similar to L2 in the trimeric LHCII of plants (Ruban et al., 2000).

**Figure 4.**
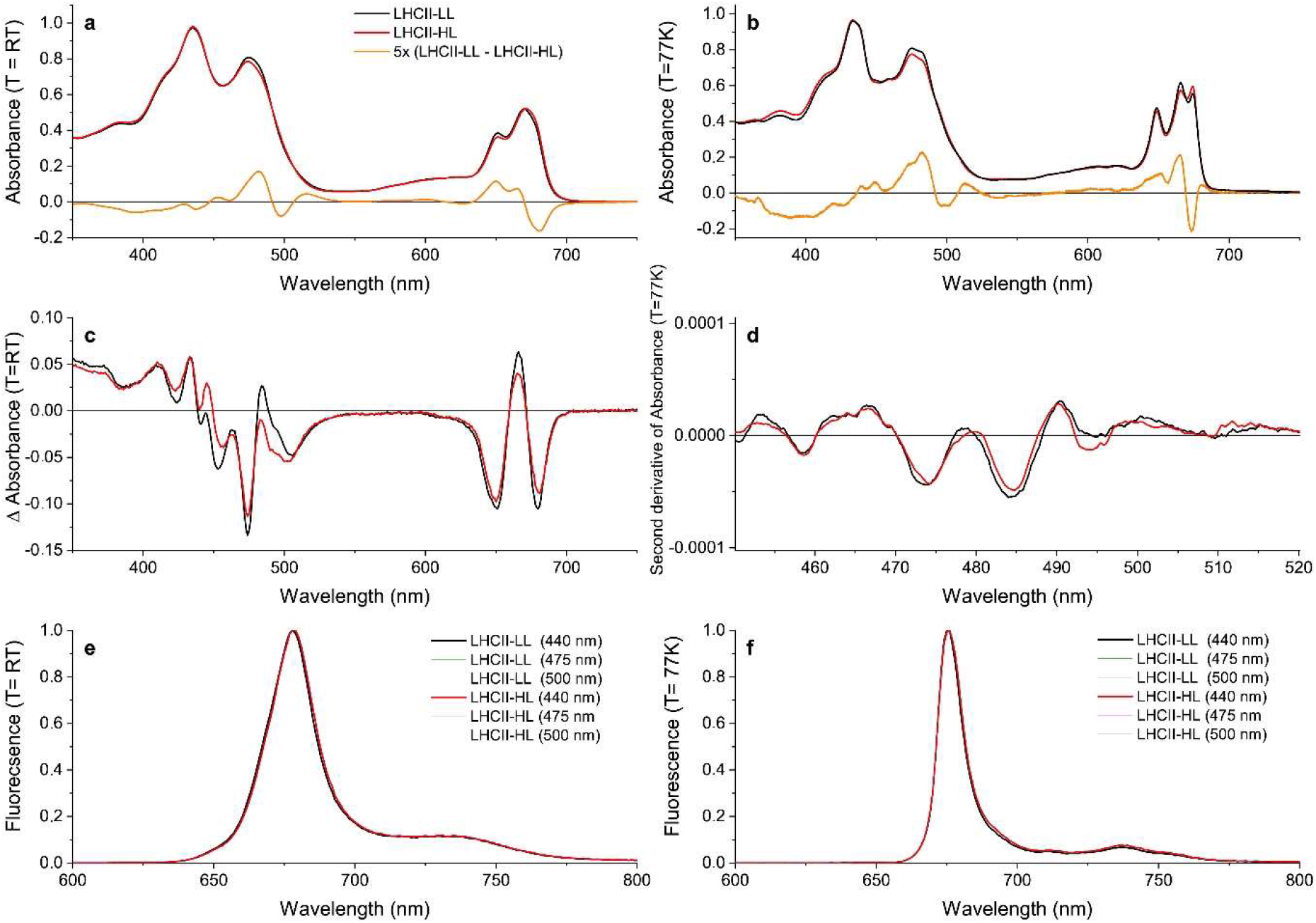
Steady-state spectra of LHCII purified from long-term LL (LHCII-LL) and HL (LHCII-HL) acclimated cells. **A** Absorption spectra at room temperature and **B** at 77K normalized to the area in the 620-700 nm region. **C** CD spectra at room temperature normalized to the respective absorption spectra **D** Savitski-golay smoothed (20 pt) second derivative of the 77K absorption spectrum. **E** Fluorescence emission spectra at room temperature and **F** at 77K. Different excitation wavelengths (440, 475, 500 nm) gave similar results for both samples. Results were reproduced on three different biological replicas or two biological replicas in the case of the 77K absorption.

### Lutein vs. Loroxanthin: different roles?

Carotenoids in LHCs are important for the stability of the complexes (Paulsen et al. 1993) and are involved in light harvesting and photoprotection by quenching excited Chl^1^ and Chl^3^ as well as scavenging singlet oxygen (Frank et al., 2004). Time-resolved fluorescence measurements demonstrate that the different pigment composition of LHCII-HL and LHCII-LL does not affect their fluorescence lifetime (Table 3), indicating that the different content of Lutein and Loroxanthin does not influence the ^1^Chl quenching, at least *in vitro*. Carotenoid to Chl EET efficiency of LHCII-HL and LHCII-LL was tested by comparing the (1-T) spectrum with the fluorescence excitation spectrum. LHCII-LL has 5 ± 1% higher Car->Chl EET efficiency than trimeric LHCII-HL (Fig. **5 (A**,**B)**, Table 3). A higher Car->Chl EET efficiency (10%) was also found for reconstituted LhcbM1 when compared with LhcbM1 reconstituted without Loroxanthin from an earlier work (Natali and Croce, 2015) (Supporting information Fig S3 and S5)

**Figure 5.**
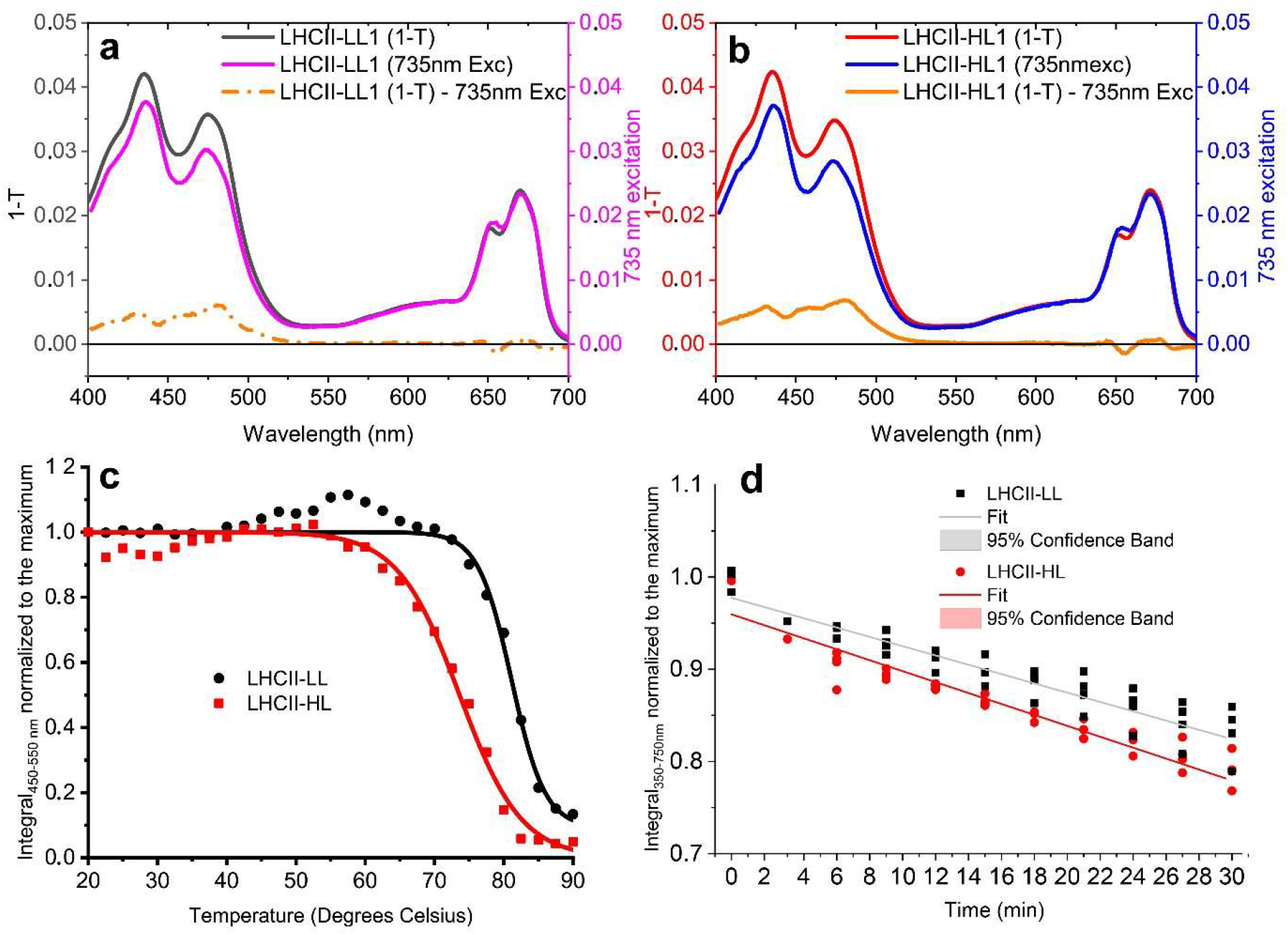
Properties of LHCII-LL and LHCII-HL. **A** (LHCII-LL) and **B** (LHCII-HL) Carotenoid to Chl EET efficiency, tested by comparing the fitted 1-T and fluorescence excitation spectra normalized to the Chl *a* content. (Full details in Supporting information Fig. S3-S4) **C**. Thermostability of LHCII-LL and LHCII-HL measured by the loss of the CD signal between 450 and 550 nm (the absolute area was used) **D**. Photo-stability of LHCII-LL and LHCII-HL tested by the photobleaching with a 4500 µmol photons m^-2^ s^-1^ halogen lamp. Data from 2 biological replicas normalized to the average integral of the initial (maximum) absorption and fitted with a linear function.

To investigate if the presence of Loroxanthin instead of Lutein influences the stability of LHCII, we followed the thermally-induced unfolding of LHCII-HL and -LL via CD measurements (Fig. 5B). Additionally, we performed photostability experiments by following the bleaching kinetics of the LHCII absorption under high-light (4500 µmol photons m^-2^ s^-1^) (Fig. 5(**C**)). The results show that LHCII-LL has a higher temperature- and photostability (Table 3) than LHCII-HL.

## Discussion

Photoacclimation mechanisms are critical for the success of photosynthetic organisms and the xanthophyll cycles have proven to be major contributors to photoacclimation for many organisms. Thus far six xanthophyll cycles have been described but more may be present, especially among algae (García-Plazaola et al., 2007). Here we presented a new XC present in *Chlamydomonas reinhardtii*: the Loroxanthin cycle. We show its kinetics and its effects on the carotenoid composition of the antenna complexes (LhcbMs). The properties of the cycle are compared to those of the other XCs and are discussed in the context of its possible physiological role.

### Changes in the Lutein hydroxylation state is a long term acclimation response in C. reinhardtii

The Lo/(L+Lo) ratio changes slowly during a “simulated summer” day, following changes in light intensity. However, the amplitude of the change of the Lutein hydroxylation state is less than half of that observed after a sustained (72hrs) change in light intensity (Fig. **3(C)**, Fig 2(**E**)). Because the change of the Lutein hydroxylation state is slow in all the conditions tested, we conclude that the Loroxanthin cycle is a long-term acclimation process. This conclusion is in agreement with other data in *C. reinhardtii* (Pineau *et al*., 2001), showing that short term high light treatment had no significant effect on the Lo/(L+Lo) ratio, and *S. Obliquus* (Bishop *et al*., 1989), showing that the Lo/(L+Lo) ratio changes by a factor of 4 upon a change in light intensity and was reversible in 48 hours.

### Loroxanthin binding sites in LHCII

The pigment content of LhcbM trimers acclimated to HL or LL (Table 2) demonstrates that a change in the Lutein hydroxylation state is also reflected in the carotenoid composition of the LHCII trimers. The LhcbM of LL grown *C. reinhardtii* bind less Lutein than the Lhcb of plants and bind Loroxanthin instead, with small stoichiometrical differences among the LhcbMs (Natali and Croce, 2015). A change in the ratio between the LhcbMs would therefore not explain the change in the carotenoid composition of the trimers from low and high light acclimated cultures. Moreover, no large changes in LHCII composition were observed previously (Bonente et al., 2012).

**Table 2.** Properties of LHCII-HL and LHCII-LL. **Fluorescence lifetime** (438 nm excitation, 680 nm detection, n=4) Errors represent the standard deviation. **Carotenoid to Chl EET efficiency**, normalized to Chl *a* (assuming 100% transfer, n=3) Errors represent the standard error (Fig. 5 (**A**,**B**), Supporting information Fig S4). **Thermostability** (Fig. 5(**B**)). **Photostability** (Fig. 5(**C**), n=2). Errors represent the error of the fit. n = number of biological replicas. **\* P < 0.05**

| LHCII | Average fluorescence lifetime (ns) | Carotenoid to Chl EET efficiency % | Thermostability transition temperature (°C) | Photostability Photobleaching rate ($\Delta\% \text{Absorption}_{350-750} \text{ nm min}^{-1}$) |
| --- | --- | --- | --- | --- |
| -LL | 3.0 ± 0.2 | 90 ± 1 * | 81 ± 0.3 | -0.50 ± 0.03 * |
| -HL | 3.0 ± 0.3 | 86 ± 2 * | 74 ± 0.3 | -0.59 ± 0.03 * |

The presence of 1.3 molecules of Loroxanthin per monomer in LHCII-LL indicates that this xanthophyll binds in at least two carotenoid binding sites. Out of the four carotenoid sites of the LhcbMs (Fig. 1), Lo was suggested to bind to L1 because its presence influences the lowest energy Chls, known to be close to L1 (Natali and Croce, 2015). This effect is also visible in the complexes analyzed here (Fig. **4(B**,**D)**). The second binding site accommodating Loroxanthin is most probably L2, since in LHCII-HL Loroxanthin is almost entirely substituted by Lutein, and it is known that out of the remaining three binding sites (L2, N1 and V1) Lutein has the highest affinity for L2 (Croce et al., 1999). Moreover, a series of experiments seems to exclude that Loroxanthin is associated with N1 and V1 in a high amount. The purification of LHC complexes via isoelectrofocusing largely removes the carotenoid bound in the V1 site while it does not affect the carotenoids in L1 and L2 (Caffarri et al., 2001; Natali and Croce, 2015). Trimeric LHCII-LL isolated with this technique lose a large fraction of Violaxanthin and Lutein and much less Loroxanthin (Natali and Croce, 2015). Similar results were observed with native PAGE of solubilized thylakoids (Pineau et al., 2001). The N1 site of plant LHCII is highly specific for Neoxanthin and this is primarily due to the presence of a tyrosine forming an H-bond with the –OH of the Neoxanthin (Caffarri et al. 2007). This tyrosine is conserved in all LhcbMs suggesting that the N1 site is specific to Neoxanthin also in these complexes. The fact that in LHCII-LL 0.8 Neoxanthin molecules are present supports this suggestion.

### Mechanistic considerations of the light-dependent change in Lutein and Loroxanthin content

Since Lutein in the L1 and L2 sites of LHCII is strongly bound to the complex, it is unlikely that an exchange of xanthophylls can occur in the folded complex. Also, the OH group of Loroxanthin is deeply buried in the transmembrane domain of LHCII and is thus unreachable from the outside. Thus, Loroxanthin and Lutein are most likely inserted in newly synthesized LHCII proteins during folding. In addition, the presence of only minimal quantities of Loroxanthin “free” in the membrane compared with Lutein and Violaxanthin (Pineau et al., 2001), suggests that the synthesis of Loroxanthin from Lutein only takes place together with the folding of new LhcbM proteins (Grossman et al., 2004).

### Role of the the light-dependent change in Lutein and Loroxanthin content

The fact that the Lutein and Loroxanthin content in cells and LhcbM is related to light intensity changes suggests opposite roles for these two xanthophylls in light-harvesting and photoprotection. The excitation energy transfer efficiency of the carotenoids associated with LHCII is higher with Loroxanthin than Lutein (5±1%). This increase is similar to that observed for Lutein-epoxide versus Lutein (Matsubara et al., 2007) and increased excitation energy transfer efficiency of the carotenoids can lead to a growth advantage in low-light environments as observed for purple bacteria (Magdaong et al., 2014). On the other hand, Lutein seems to be important in high light conditions. One of the possibilities is that Lutein is a better Chl^3^ quencher or oxygen scavenger than Loroxanthin. However, this does not seem to be the case since the photostability of LHCII-LL is high and even slightly higher than that of LHCII-HL, meaning that both complexes are well protected. The thermo-stability of LHCII-LL is also increased compared to LHCII-HL and both complexes remain completely stable up to 60 °C (Fig. **5(C)**), which is above the physiological temperature range of *Chlamydomonas reinhardtii* (Starks *et al*., 1981). The additional incorporation of Lutein in the LhcbM in HL might be important for NPQ. The availability of mutants of the Lutein hydroxylation enzyme would allow testing this hypothesis. The fact that the lifetime of LHCII *in vitro* does not change in the presence of Lutein or Loroxanthin is not conclusive in this respect because it is known that isolated LHCII are stable in their light-harvesting conformation (Horton et al., 1996). However, the observation that NPQ can occur in the absence of Lutein and Loroxanthin (Niyogi et al., 1997) and even when non-native xanthophylls are associated with LHCII (Xu et al., 2020) suggests that the Lutein to Loroxanthin exchange might not have a significant effect on quenching in individual LHCs. Alternatively, the additional incorporation of Lutein in the LhcbM in HL could be a side-effect of the increase of Lutein content in the membrane upon high-light acclimation (Bonente et al., 2012; Polukhina et al., 2016) that may provide an increase in photoprotection that overcomes the disadvantages of additional Lutein binding to the LhcbMs.

### Comparison of the Loroxanthin cycle with other xanthophyll cycles

The Loroxanthin cycle has some similarity with the LLx cycle of plants: (1) Lutein-epoxide has increased EET to Chl compared with Lutein (+7.9%) and mostly binds to the Internal Lutein binding sites of LHCII (Matsubara et al., 2007). (2) Complete LL Lo/L and Lo/100 Chl (*a*/*b*) ratios take more than 24 hours to be attained, similar to the truncated LLx cycle (Esteban and García-Plazaola, 2014). However, there are also differences: (1) hydroxylation versus epoxidation of Lutein; (2) The Loroxanthin cell content of LL grown *Chlamydomonas reinhardtii* is higher than the Lutein-epoxide content in more than 95% of the species that contain the LLx cycle (Esteban and García-Plazaola, 2014) (3) the Loroxanthin content of LhcbM of *C. reinhardtii* (Lo/L (1)) is higher than the Lutein-epoxide content of the Lhcb of shade plants (Lx/L 0.47-0.8 (Matsubara et al., 2003, 2005; García-Plazaola et al., 2007)).

The Loroxanthin cycle differs from the VAZ, in two main aspects: it is far slower and it leads to a change in the occupancy of the L1/L2 binding sites of LHCII, which is not the case for VAZ (Xu et al., 2015). Interestingly, the presence of both cycles in *C. reinhardtii* suggests that they are involved in slow and fast photoprotection strategies. A xanthophyll cycle operating at longer timescales than the VDE cycle, such as the LLx cycle (García-Plazaola et al., 2007; Esteban and García-Plazaola, 2014) and the Loroxanthin cycle, is thus likely to provide evolutionary advantages for photosynthetic organisms that experience long periods of low light. Loroxanthin is present in algae of the Chlorophyte, Euglenophyte and Chlorarachniophyte (Takaichi, 2011) and in addition to *C. reinhardtii* its content has been shown to fluctuate with the light intensity in the *Chlorophytes Botryococcus braunii* (van den Berg et al., 2019), *Tetraselmis suecica* (Garrido et al., 2009) *and Scenedesmus obliquus* (Senger et al., 1993). The loroxanthin cycle is therefore likely widespread and possibly active in all algae that contain Loroxanthin, similar to the LLx cycle in Lutein-epoxide containing plants (García-Plazaola et al., 2007).

## Supplementary Figures

**Fig. S1 Time course of cellular Chl content (mg/mL) (A). Chl/Car ratio (mol mol** ^**-1**^**) (B) and Chl *a*/*b* ratio (mol mol** ^**-1**^**) (C) of the five biological replicas following the shift from HL to LL**.

**Table S1 Students paired one-tailed t-test results of the different timepoints in Fig. 3 (B,C)**

**Fig. S2 Changes in *C. reinhardtii* cell culture during a “summer” day (18:6 (D:N) sinusoidal light regime)**

**Fig. S3 Example of the Fitting of 1-T and Fluorescence excitation spectra to determine Car - > Chl *a* Excitation Energy Transfer efficiency**

**Fig. S4 Efficiency of Excitation energy transfer (EET) of LHCII-LL (three biological replicas A**,**B**,**C) and LHCII-HL (three biological replicas d**,**e**,**f) calculated by comparing the 1-T spectrum with the fluorescence excitation spectrum**.

**Fig. S5 Efficiency of Excitation energy transfer (EET) of LhcbM1 (A) and LhcbM1 – No Loroxanthin (B)** (Natali and Croce, 2015) **calculated by comparing the 1-T spectrum with the fluorescence excitation spectrum**.

## Acknowledgments

Dr. Bart van Oort is acknowledged for helpful discussions.

## Author contributions

T E van den Berg conceived the research, T E van den Berg performed the experiments, T E van den Berg and R Croce analyzed the data. T E van den Berg and R. Croce wrote the manuscript.

## Data availability

Data is available upon reasonable request

## Notes

### Competing Interest Statement

The authors have declared no competing interest.

