## Supplementory information for "The Loroxanthin cycle: A new type of xanthophyll cycle in green algae (Chlorophyta)"

The following Supporting Information is available for this article:

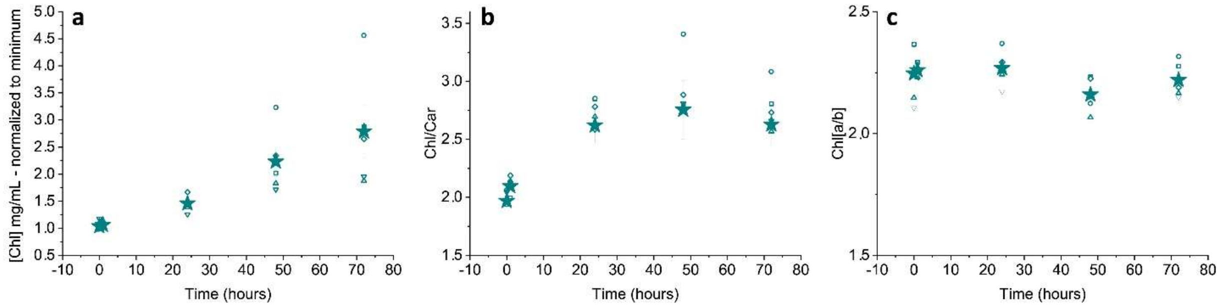


**Fig. S1 Time course of cellular Chl content (mg/mL) (A). Chl/Car ratio (mol / mol) (B) and Chl *a*/*b* ratio (mol / mol) (C) of the five biological replicas following the shift from HL to LL.** Chl/Car is significantly different from 0hrs (HL) from 1 hour LL onwards (P<0.05). Four technical replicas per biological replica. Star symbols represent the average. The error bar represents the standard error of the mean.

**Table S1 Students paired one-tailed t-test results of the different timepoints in figure 3.** Lo/(L+Lo) Fig. 3C **(B).** ((0.5*A)+Z)/(V+A+Z) Fig. 3B P values below 0.05 level are indicated by a black number and above by a red number.

| A | t(0) | t(3) | t(6) | t(9) | t(12) | t(15) | t(18) |
|---|---|---|---|---|---|---|---|
| t(3) | 0.19 | | | | | | |
| t(6) | 0.43 | 0.01 | | | | | |
| t(9) | 0.12 | 0.01 | 0.01 | | | | |
| t(12) | 0.08 | 0.01 | 0.01 | 0.05 | | | |

| | | | | | | |
|---|---|---|---|---|---|---|
| t(15) | 0.48 | 0.02 | 0.34 | 0.04 | 0.04 | |
| t(18) | | | | | | |
| t(24) | 0.14 | 0.15 | 0.06 | 0.03 | 0.03 | 0.04 |

| **B** | **t(0)** | **t(3)** | **t(6)** | **t(9)** | **t(12)** | **t(15)** | **t(18)** |
|---|---|---|---|---|---|---|---|
| t(3) | 0.01 | | | | | | |
| t(6) | 0.05 | 0.08 | | | | | |
| t(9) | 0.02 | 0.02 | 0.16 | | | | |
| t(12) | 0.03 | 0.10 | 0.08 | 0.00 | | | |
| t(15) | 0.03 | 0.01 | 0.01 | 0.00 | 0.00 | | |
| t(18) | 0.24 | 0.02 | 0.03 | 0.00 | 0.00 | 0.02 | |
| t(24) | 0.28 | 0.01 | 0.00 | 0.00 | 0.01 | 0.01 | 0.46 |

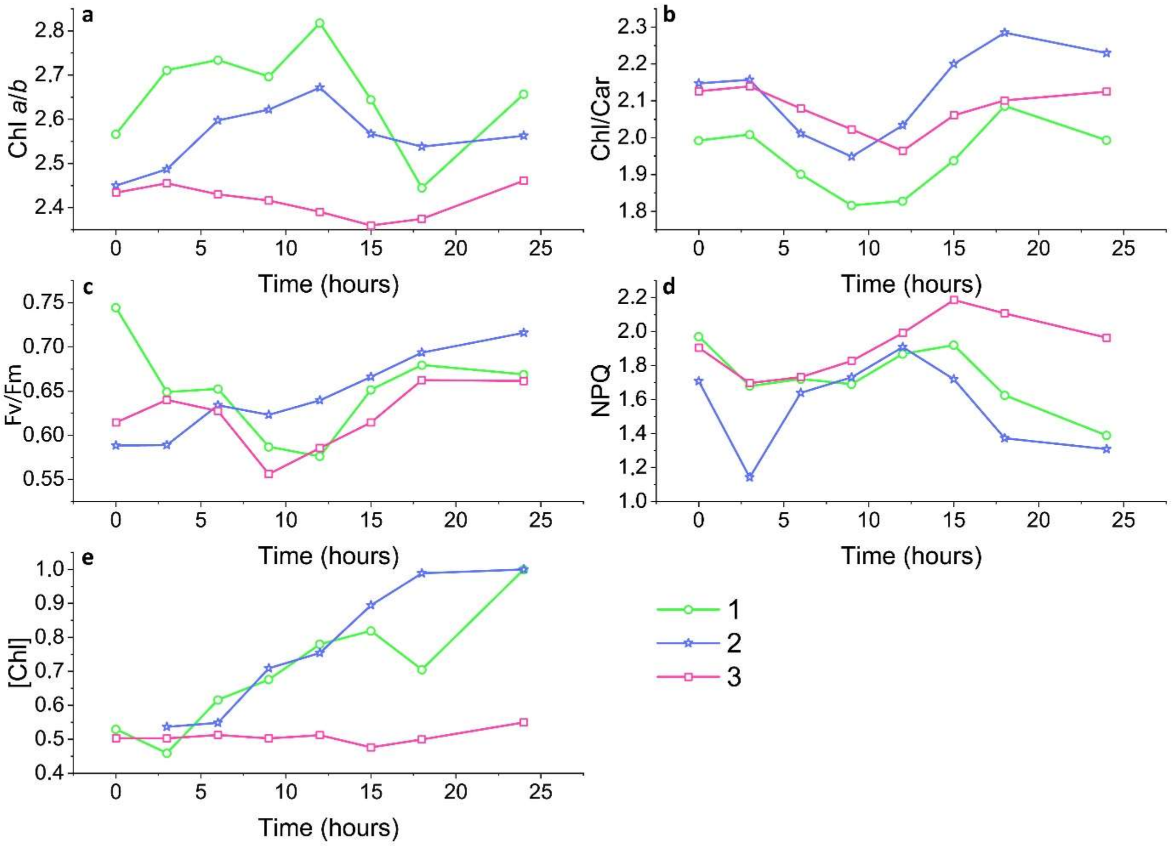


**Fig. S2 Changes in C. reinhardtii cell culture during a "simulated summer" day (18:6 (D:N) sinusoidal light regime)** (**A**) Chl *a*/*b* (mol / mol) (**B**), Chl/car (mol / mol) (**C,D**) Cells were taken from the culture and dark-adapted for twenty minutes in a quartz cuvet while stirring in a Dual-PAM. $F_m$ and $F_0$ were determined and then an induction curve was taken for 15 min with 2080 µmol photons $m^{-2}$ $s^{-1}$ actinic light. F' and $F_m$' were measured every 3 min and $F_v/F_m$ and NPQ were calculated as: $\frac{\mathrm{Fv}}{\mathrm{Fm}} = \frac{\mathrm{Fm}' - \mathrm{F}}{\mathrm{Fm}'}$ and $\mathrm{NPQ} = \frac{\mathrm{Fm} \quad \mathrm{Fm}'}{\mathrm{Fm}'}$ . $F_v/F_m$ (c) and final (after 15 min) NPQ (d) values were plotted. (**E**) [Chl] in mg/ml during 24 hours of cultivation of three biological replicas normalized tot the maximum. Biological replica 3 deviated from biological replica 1 and 2 because the Chlorophyll concentration and Chl a/b of the culture did not change during the 24 hours of measurements. We suspect that replicate three was perhaps diluted too much and therefore experiencing a lag phase in growth, leading to a lack of increase of the Chl concentration

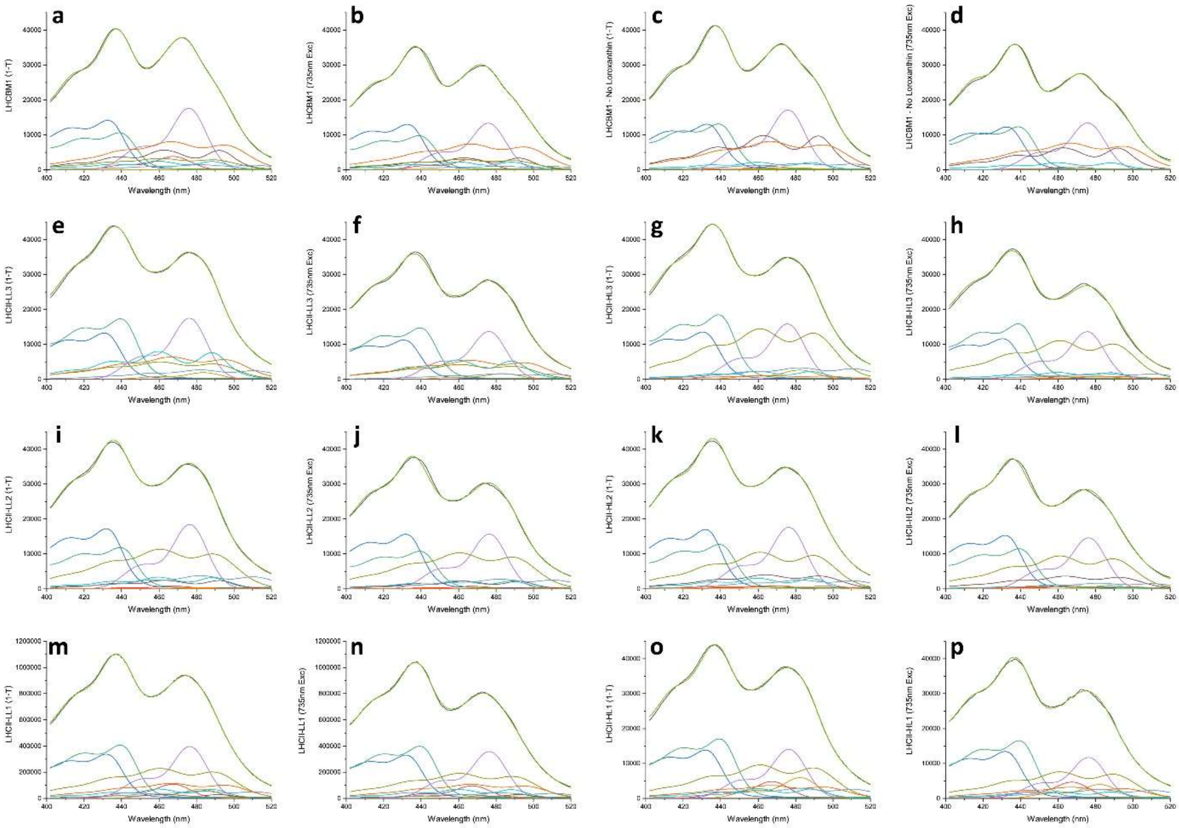


**Fig. S3 Fitting of 1-T and Fluorescence excitation spectra to determine Car -> Chl *a* Excitation Energy Transfer efficiency. (A) (LHCBM1), (C) (LHCBM1 – No loroxanthin), € (LHCII-LL3), G (LHCII-HL3), (I) (LHCII-LL2), (K) (LHCII-HL2), (M) (LHCII-LL1), and (O) (LHCII-HL1) (1-T) (B) (LHCBM1), (D) (LHCBM1 – No loroxanthin), (F) (LHCII-LL3), (H) (LHCII-HL3), (J) (LHCII-LL2), (L) (LHCII-HL2), (N) (LHCII-LL1), and (P) (LHCII-HL1) (735nm Excitation spectrum)**. The 1-T and excitation spectrum are normalized to the fitted quantity of Chl *a* in each spectrum. Spectral positions and extinction coefficients were taken from ((Croce *et al.*, 2000; Caffarri *et al.*, 2001; van den Berg *et al.*, 2018).

For LHCII-LL and LhcbM1, two Loroxanthin, one Lutein, one Violaxanthin and one Neoxanthin spectral forms and for LHCII-HL three Lutein, one Loroxanthin and one Neoxanthin spectral forms were used for the fitting. For 'LhcM1 - No Loroxanthin', two Luteins , one Violaxanthin and one Neoxanthin spectral forms were used for the fitting. The spectra used for fitting were chosen based on the pigment composition determined by hplc. LHCBM1 reconstitutions from (Natali & Croce, 2015)**.** Restrictions for the fit were: 1. Individual spectral shapes and positions are the same in all 1-T and excitation spectra. 2. Individual Chl *b* and carotenoid concentrations relative to Chl *a* are similar or lower in the excitation spectrum than in the 1-T spectrum. Carotenoid EET in Table 3 is calculated by: $\frac{\text{integral fitted carotenoid spectra LL 735nm Exc}_{(400-520\text{nm})}}{\text{integral fitted carotenoid spectra LL 1}-\text{T}_{(400-520\text{nm})}} - \frac{\text{integral fitted carotenoid spectra HL 735nm Exc}_{(400-520\text{nm})}}{\text{integral fitted carotenoid spectra HL 1}-\text{T}_{(400-520\text{nm})}} \times 100\%$

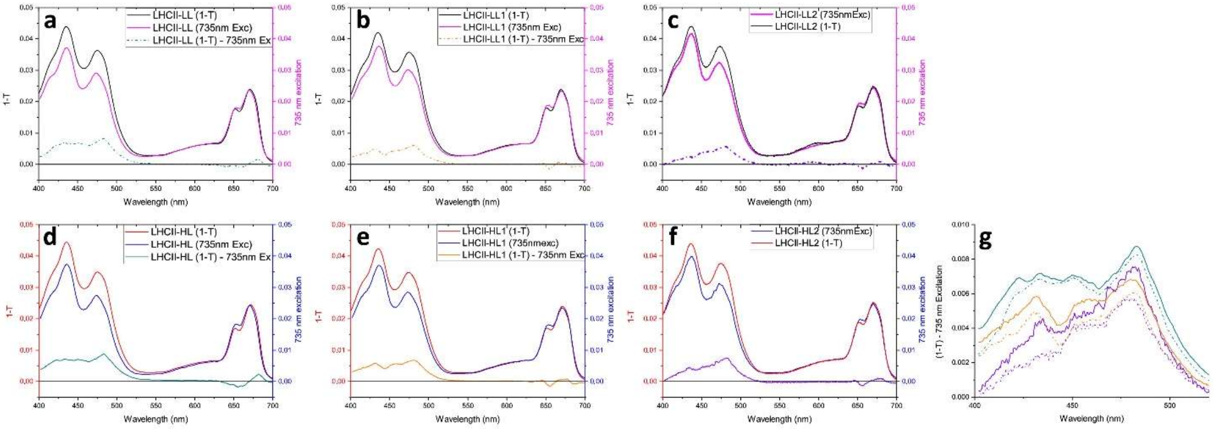


**Fig. S4 Efficiency of Excitation energy transfer (EET) of LHCII-LL (three biological replicas (A,B,C) and LHCII-HL (three biological replicas (D,E,F) calculated by comparing the 1-T spectrum with the fluorescence excitation spectrum.** Chl *a & Chl b* -> Chl *a* EET is assumed to be 100% and therefore the spectra are normalized against their integral from 620 to 700 nm. **(G)** Difference spectra of all biological replicas. Total EET difference between LHCII-LL and LHCII-HL expressed as: $\left(\frac{(LL\ 735nm\ Exc)_{integral\ 400-520nm}}{(LL\ 1-T)_{integral\ 400-520nm}}\right) - \left(\frac{(HL\ 735nm\ Exc)_{integral\ 400-520n}}{(HL\ 1-T)_{integral\ 400-520nm}}\right) \times 100\%$, is 2.8 ± 0.4% in all biological replica's (P=0,019)

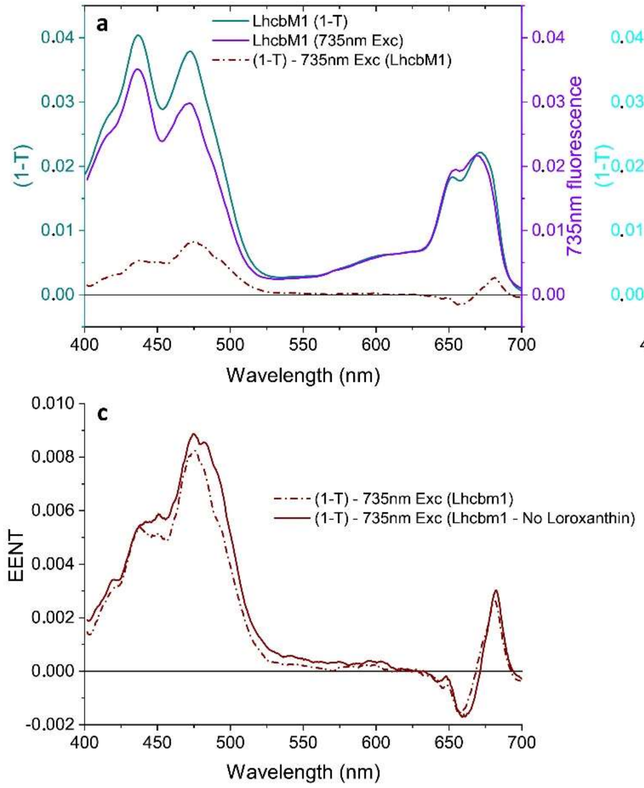


**Fig. S5 Efficiency of Excitation energy transfer (EET) of LhcbM1 (A) and LhcbM1 – No Loroxanthin (B)** (Natali & Croce, 2015) **calculated by comparing the 1-T spectrum with the fluorescence excitation spectrum.** Chl *a & Chl b* -> Chl *a* EET is assumed to be 100% and therefore the spectra are normalized against their integral from 620 to 700 nm. **(C)** Difference spectra of both samples. EENT is excitation energy not transferred. <u>Total</u> EET difference expressed as:

$$\left(\frac{(LHCB \quad 735nm\ Exc)_{integral\ 400-520nm}}{(LHCBM1\ 1{-}T)_{integral\ 400-520n}}\right) - \left(\frac{(LHCBM1{-}No\ Loroxanthi \quad 735nm\ Exc)_{integral\ 400-520nm}}{(LHCBM1{-}No\ Loroxanthi\ )_{integral\ 400-520nm}}\right) \times 100\%$$

, is 2.3%.
